# Four new threatened species of *Rinorea* (Violaceae), treelets from the forests of Cameroon

**DOI:** 10.1101/2021.05.19.444792

**Authors:** Gaston Achoundong, Xander van der Burgt, Martin Cheek

## Abstract

Four species of *Rinorea* are described as new to science; all four species are endemic to evergreen rain forest in Cameroon. *Rinorea villiersii* Achound., and *Rinorea amietii* Achound. are placed in Sect. *Crassiflorae* Wahlert, while *Rinorea dewildei* Achound. and *Rinorea faurei* Achound. fall in *Rinorea* section *Dentatae* (Engl.) Wahlert. The first species appears to be endemic to the Solé Forest Reserve northeast of Yabassi in Littoral Region. The second and the third species are found mainly in Littoral and South Regions, *Rinorea amietii* extending to South West Region and *Rinorea dewildei* extending to Central Region. The fourth species, *Rinorea faurei* is an endemic of the Santchou Forest Reserve at the foot of Dschang Plateau in West Region. The affinities of the four species are discussed, they are illustrated and mapped, and their conservation status is assessed. All four species are threatened with extinction according to the 2012 IUCN categories and criteria.

## Introduction

In the course of revising the species of Violaceae of Africa, mainly in preparation for the account of the Violaceae for the “Flore du Cameroun”, the first author has, with various collaborators, already published 13 new species to science for this group (Achoundong & Onana 1998; Achoundong & Bos 1999; Achoundong & Bos 2001; Achoundong (2003); Achoundong & Cheek 2003; Achoundong & Cheek 2005; Achoundong & Bakker 2006). He has also contributed to numerous checklists and other publications (Achoundong 1996; Achoundong 1997; Amiet & Achoundong 1996; Achoundong 2000; Bakker *et al*. 2006). These latter publications include several provisional names which have been used for the new species published in the present paper by the first author and others. In this paper, four such species names are formally published to validate these names.

The most recent phylogenetic studies and classification of African *Rinorea* Aubl. are set out by Wahlert (2010), Wahlert & Ballard (2012) and van Velzen *et al*. (2015). The first two new species described below, *Rinorea villiersii* Achound. and *Rinorea amietii* Achound. fall within *Rinorea* section *Crassiflorae* Wahlert (2010), while the second two species, *Rinorea dewildei* Achound. and *R. faurei* Achound. fall in *Rinorea* section *Dentatae* (Engl.) Wahlert.

The genus *Rinorea* is pantropical, with 206 species currently accepted by Plants of the World Online (POWO, continuously updated, accessed April 2021). Africa is the most species-diverse continent for *Rinorea* with 110-150 species (van Velzen *et al*. 2015). *Rinorea* species are mainly forest understorey shrubs or small trees. Morphologically they are characterized by having alternate, simple leaves, often with petioles of different lengths on the same stem and a long, curving apical bud. The flowers are often green, dull yellow, or shades of white and are usually zygomorphic. There are three sets of petals in *Rinorea:* an anterior petal (also known as the lower or ventral petal), two lateral petals and two posterior petals. These are likely homologous to the three sets of petals in other strongly zygomorphic genera of Violaceae, such as *Viola* L. (Wahlert 2010).

The anterior petal is larger than the other petals, and often modified, with taxonomically important, often diagnostic characters. The androecium has a staminal tube which is also zygomorphic: the anterior (lower or ventral) side is longer and entire, while on the dorsal side the staminal tube is generally shorter and incised with a V-shaped cleft.

The fruits are typical of the family: dry, tricoccal, septicidal capsules with parietal placentation. *Rinorea* are ecologically important and diverse in African forests, often with several sympatric species in one forest, for example ten species were recorded in a few square kilometers of the Mefou Proposed National Park near Yaoundé (Cheek *et al*. 2011). Many species are very range-restricted, found in such small areas that they are at risk of extinction from forest clearance. *Rinorea dewitii* Achound., *R. fausteana* Achound., *R. simoneae* Achound. and *R. thomasii* Achound. are all assessed as threatened in the Red Data Book of the Flowering Plants of Cameroon (Onana & Cheek 2011) and all but the first can be found on the IUCN Red List (iucnredlist.org) e.g., *Rinorea thomasii* (Darbyshire & Cheek 2004a; Cheek 2017; Darbyshire & Cheek 2014b). In neighbouring Gabon, the recently published *Rinorea calcicola* Velzen & Wieringa is also range-restricted and of conservation concern (van Velzen & Wieringa 2014). Cameroon has the highest species-diversity for the genus in tropical Africa with 53 species listed (Onana 2011), followed by Gabon, with 46 species (Sosef *et al*. 2006). However, the superficial similarity between species has made identification difficult for taxonomists, e.g., 194 specimens of *Rinorea* are listed as unidentified to species for Gabon in Sosef *et al*. (2006).

African *Rinorea* species are of great interest to entomologists, being important larval food plants of the butterfly genus *Cymothoe*. Twenty-eight species of *Cymothoe* are known to feed on *Rinorea*, of which 18 are strictly monophagous, six are oligophagous and three feed on up to six species of *Rinorea* (Amiet & Achoundong, 1996).

## Materials & Methods

All specimens cited have been seen. Herbarium citations follow Index Herbariorum (Thiers *et al*. continuously updated). Specimens were studied online, on loan from, or at BR, K, P, WAG and YA. We also searched JSTOR Global Plants (https://plants.jstor.org/ accessed April 2021) for additional materials. Binomial authorities follow the International Plant Names Index (IPNI 2021), and nomenclature follows Turland *et al*. (2018). The conservation assessment was made using Bachman *et al*. (2011) following the categories and criteria of IUCN (2012). Herbarium material was examined with a Leica Wild M8 dissecting binocular microscope fitted with an eyepiece graticule. Measurements are made from rehydrated material. The terms and format of the description follow the conventions of Achoundong & Cheek (2005). Georeferences for specimens were obtained from Google Earth. (https://www.google.com/intl/en_uk/earth/versions/).

## Results: Taxonomy

> **1. *Rinorea villiersii*** *Achound*. **sp**.**nov**. Type: Cameroon, Littoral Region, Sole, north-west of Yabassi, fl. 20 May 1992, *Achoundong* 1928, (holotype YA; isotypes, P, WAG)

*Rinorea villiersii* Achound. ined. (Achoundong 1996, 1997; Onana 2011: 151)

*Rinorea solensis* Achound. ined. (Achoundong 1996, 1997).

*Evergreen shrub or treelet* 1-2 (−3) m tall, densely branched. Stems and leaves pubescent when young, glabrous at maturity. *Leaves* white when young, subcoriaceous, elliptic, 11-20 (−22) x 3-5(−7) cm strongly long acuminate, base cuneate, margin serrate; lateral nerves 10-13 on each side of the midrib. Petiole 1.5-2(−3) cm long. Stipules triangular, 2-3 (−4) x 0.5-1(−2) mm, mucronate, with two prominent longitudinal nerves, margins crenate, ciliate. *Inflorescence* terminal, corymbiform, 2.5-3.5 (−5) x 1.5-2 (−3) cm.

Bracts 5-6, triangular, 3-4(−5) x 1-2 (−3) mm, mucronate, ciliolate, pubescent. *Flowers* zygomorphic, 3-4(−5) x 2-3 (−4) mm; sepals 5, green, triangular, (1-2(−3) x 0.5-1(−1.5) mm ciliolate. *Petals* 5, white, ovate, 4-5 (−6) x 1-1.5(−2) mm; lower petal broader than the upper and the lateral petals, constricted towards the apex, inner surface densely hairy towards the apex. Androecium 3-4 mm long; staminal tube 0.5-1(−1.5) mm long, margin entire, undulate, with a V-shaped incision in the dorsal face; filaments inserted on the inner face of staminal tube; staminal tube border hiding the lower part of anthers, anthers 0.5-1(−1.5) mm long; connective appendage red, ovate, up to 0.5-1(−1.5) mm long, thecal appendages filiform, bifid. Ovary fusiform, hairy, 1-1.5(−2) mm long; style erect, 1-1.5 (−2) mm long. *Fruit* white, tricoccal, fusiform 1-1.5(−2) x 1-1.5 cm, tuberculate. *Seeds* six per fruit white, tetrahedric. Fig 1

**Fig. 1.**
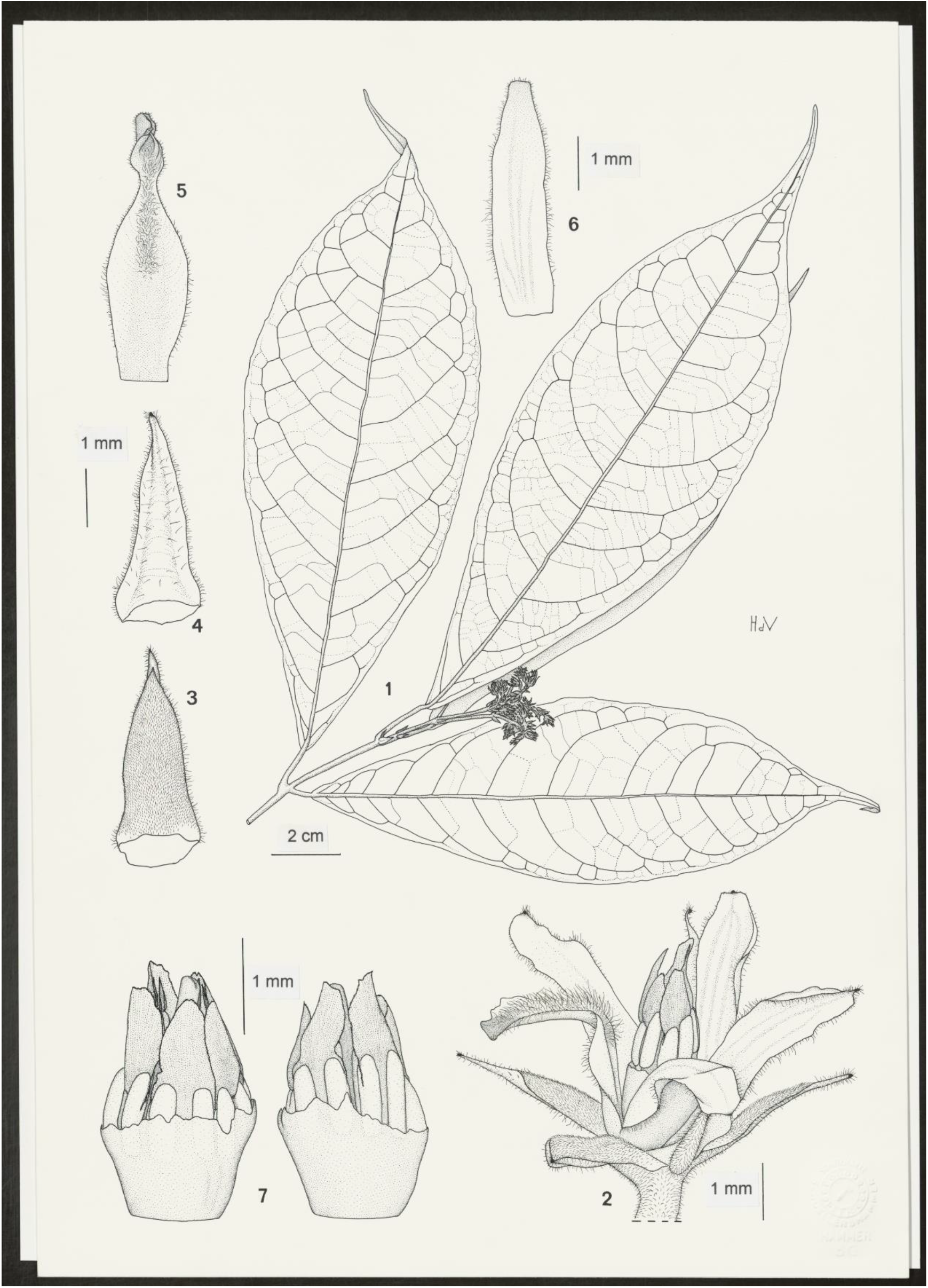
*Rinorea villiersi* **1** habit, flowering stem; **2** flower, side view; **3** sepal adaxial view; **4** sepal abaxial view; **5** dorsal (large) petal, adaxial view; **6** non-dorsal (one of four smaller) petal, adaxial view; **7** staminal tube, side views. **1 & 2** from *Achoundong* 2212 (WAG); **3-7** from *Achoundong* 1928 (WAG). Drawn by J.M. (HANS) DE VRIES

### RECOGNITION

*Rinorea villiersii* Achound. is similar to *Rinorea umbricola* Engl., in habit, the long sepals, and the large androecium. It differs in the white juvenile leaves (vs green), the leaves long-acuminate with cuneate bases, pubescent, lacking gland dots on the lower surface (vs acuminate, cordate, glabrous, with gland dots on the lower surface), the ovary fusiform, pubescent (not ovoid, glabrous). The two species can be separated by the following key characters:

Lamina 10 – 12 × 3.5 – 4.5 cm, glandular beneath; ovary glabrous *Rinorea umbricola*

Lamina 11 – 22 × 3 – 7 cm, not glandular beneath; ovary pubescent *Rinorea villiersii*

### DISTRIBUTION

Cameroon. *Rinorea villiersii* is endemic to the Solé Forest, Solé - Dibong area, north west of Yabassi town, Littoral Region (Map 1).

### SPECIMENS STUDIED. CAMEROON

**Littoral Region**, Solé, Yabassi area, fl., 15 March 1991, Amiet in *Achoundong* 1792 (P, YA); ibid., fl. 20 May 1992, *Achoundong* 1928 (holotype YA; isotypes P, WAG); ibid., fl. fr., May 1994, *Achoundong* 2212 (WAG, YA); Yabassi area, river bank in Solé Forest Reserve, fl., 9 May 2004, *Achoundong* 2328 (YA).

### HABITAT

*Rinorea villiersii* is known from lowland evergreen rain forest in the Solé-Dibong area northwest of Yabassi. The relief in its range is undulating, with valleys and plains. All specimens were collected from the borders of a river, at altitudes from 20 to 100 m.

### CONSERVATION STATUS

*Rinorea villiersii* is only known from the Solé Forest Reserve. On current evidence, the observations of the first author, we estimate the total extent of occurrence of *Rinorea villiersii*, equating to the global area of occupation, as 20 km^2^. This forest currently has good conservation status in that much of it is intact (Google Earth imagery viewed 11 April 2021) but there are no concrete conservation actions in place. The area has a very low human population density. Unfortunately, it is only 14 km along a major road from the settlement of Loum which is a very highly populated area, and which has active farmers. They are expected in the future to invade the reserve for farming purposes, resulting in major threats to this species.

Botanical surveys and other plant studies for conservation management in forest areas north, west and east of Solé Forest Reserve have resulted in many thousands of specimens being collected and identified, but failed to find any additional specimens of *Rinorea villiersii* (Cable & Cheek 1998; Cheek *et al*. 2000; Maisels *et al*. 2000, Chapman & Chapman 2001; Harvey *et al*. 2004; Cheek *et al*. 2004; Cheek *et al*. 2010; Harvey *et al*. 2010; Cheek *et al*. 2011). Although there are still poorly sampled locations with intact natural habitat in Cameroon, it is possible that *Rinorea villiersii* is truly localised, range-restricted and threatened as are several other species of the genus (see introduction). *Rinorea villiersii* is therefore here assessed, on the basis of the single location, small known range-size and threats described above, as Critically Endangered, CR B1+2ab(iii).

### ETYMOLOGY

The specific epithet commemorates the late Dr Jean François Villiers (1943-2001), taxonomic specialist in mimosoid Leguminosae (also known as Mimosaceae) of Africa and Madagascar, staff member of the Herbarium at MNHN, Paris and lecturer at the University of Yaoundé. He was the supervisor of the first author’s doctoral thesis on *Rinorea*. He did much to encourage botanists in Cameroon to study conservation.

### VERNACULAR NAMES & USES

None are recorded.

> **2. *Rinorea amietii*** *Achound*. **sp**.**nov**. Type: Cameroon, Littoral Region, Marienberg, March 1992, fl. *Achoundong* 1853 (holotype P; isotypes WAG, YA).

*Rinorea amietii* ined. Achoundong (1996: 544); Amiet & Achoundong (1996: 465); Achoundong (1997); Onana (2011: 150)

*Shrub or small tree* 5-7 (−8) m high. Diameter of the trunk at 1.3 m above the ground up to 10 cm. Stems and leaves ferruginously hairy, most densely pubescent when young. Leaves subcoriaceous, elliptic, 9-17(−20) x 3-7(−9) cm, broadly acuminate, base cuneate, margin serrate, lateral nerves 12-13 on each side of the midrib. Petiole 5-3(−4) cm long. Stipules triangular, 2-5 × 1-1.5 mm, mucronate, densely pubescent.

*Inflorescence* terminal or subterminal, pubescent, corymbiform, 1.5-5 (−6) x 1.5-2.5 (−4) cm, bracts triangular 3-5(−6) x 1-1.5 (−2) mm, pubescent, densely so on the midrib, apiculate. *Flower* zygomorphic, 4-5(−6) x 3-4 mm. Sepals 5, bright yellow, oblong 1.5-2.5 (−3) x 0.5-1.5(−2) mm, pubescent. Petals 5, yellow, ovate to elliptic, 2-4 (−5) x 1-1.5(−2) mm, narrowed at the base, apex rounded, ciliate. Lower petal enlarged at the middle, narrowed at the extremities, more strongly near the apex; hairy at the apex of the inner face. *Androecium* 2.5-3.5(−4) mm long; staminal tube up to 1-1.5 (−2) mm high; margin free, undulate, with a V-shaped incision in dorsal face; filaments all inserted on the inner face of the staminal tube; anthers 0.5-1.5 (−2) mm long; connective appendage red, ovate 1-1.5 (−2) mm long, thecal appendages bifid, filiform. Ovary bottle-shaped, 0.5-1(−1.5) mm long, glabrous; style erect, 1.2-2 (−2.5) mm long, enlarged at the summit, with longitudinal furrows. *Fruit* ovoid, 2 × 1.5 cm, surface white, slightly tuberculate. *Seeds* six, black, angular, dorsal faces convex, inner faces plane. Fig 2

**Fig. 2.**
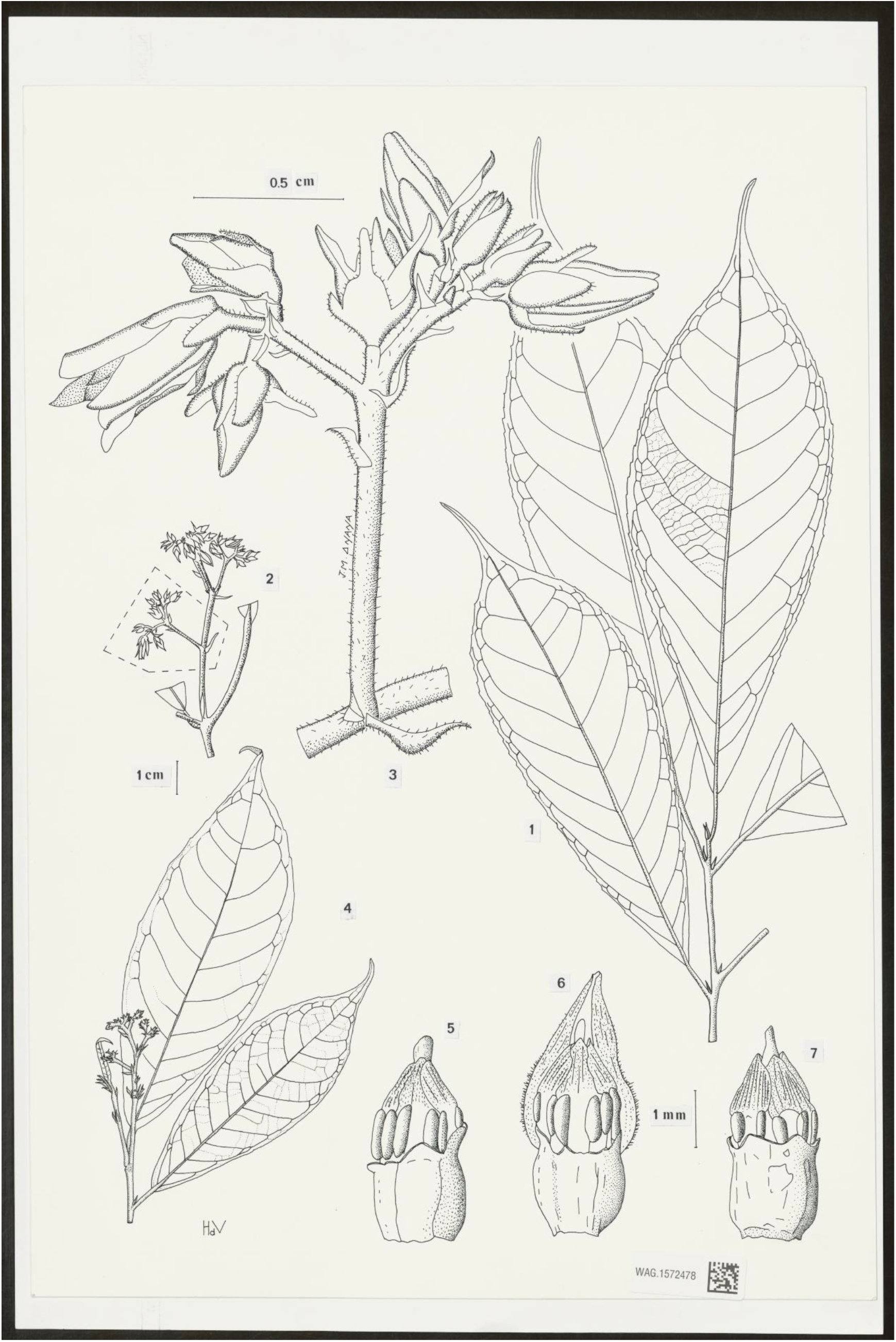
*Rinorea amietii* **1** habit, sterile leafy shoot; **2** habit showing position of terminal inflorescence; **3** detail of 2 showing portion of inflorescence; **4** habit, stem with inflorescence and buds; **5-7** views of staminal tube. **1-3 & 4-7** from *Achoundong* 1744 (YA)**; D** from *Achoundong* 1853 (WAG). **A-C & E-G** drawn by JEAN MICHEL ONANA, **D** by J.M. (HANS) DE VRIES.

### RECOGNITION

In its habit, its ferruginous leaves and its very short apical stem bud, *Rinorea amietii* is morphologically close to *Rinorea subsessilis* Brandt. The two species are separated by following characters:

> Lamina glabrous, glandular below, fruit green, smooth, inflorescence glabrous, 3-15 cm long, sepals glabrous, ovate, apex rounded not mucronate, petal midrib glabrous ............................................................................................................ *R. subsessilis*
>
> Lamina pubescent, not glandular below, fruit white, verrucose, inflorescence pubescent, 2-5 cm long, sepals densely pubescent, triangular to trapezoid, acute at the summit, mucronate, petal midrib pubescent ............................................................ *R. amietii*

## DISTRIBUTION

Cameroon. *Rinorea amietii* is endemic to coastal (littoral) Cameroon in the Southwest Region, Littoral Region and South Region (Map 1).

### SPECIMENS STUDIED. CAMEROON

**Littoral Region**, Dizangue area, Ossa Lake, fl. 5 April 1984, *Achoundong* 944 (P, YA); km 30, Edéa-Douala road, fl. 13 Jan. 1990, *Achoundong* 1550 (P, YA); Mangonbe border, fl. 2 March 1990, *Achoundong* 1627 (K, P, YA); 30 km on Edéa-Douala road, fl., fr., 28 July 1990, *Achoundong* 1744 (WAG, YA); Bonépoupa, 30 km on Edéa-Douala road, fl. 22 May 1991, *Achoundong* 1824 (P, YA); km 30 Edéa-Bonépoupa road, fl. 22 May 1991, fl., fr. *Achoundong* 1836 (P, YA); Mangonbé border, Edéa-Songloulou road, 22 May 1991, fl. *Achoundong* 1839 (YA); Littoral Region, Marienberg, March 1992, fl. *Achoundong* 1853 (holotype P; isotype, K); region de Mouanko, fl. March 1992, *Achoundong* 1864 (YA); Kribi area, Bidou I, fl. March 1993, *Achoundong* 2011 (YA); Bipindi area, Nsola, fl., March 1993, *Achoundong* 2020 *bis* (YA); Bipindi area, Nsola, fl., June 1994, *Achoundong* 2204 (YA); Edea-Douala road 19 Feb. 2001, *Achoundong* 2106 (YA); 20 km from Douala on Edea road, Carriere Ducam Duclair, 20 Dec.2002, *Achoundong* 2198 (YA); 25km NW of Edéa, SW of “Carrière Ducam Duclair”, forest at field of Marcel Melingui Engama 3° 55.01’N, 10° 03.30’E, fr. 5 May 2006 *van Velzen* 50 (WAG041636); 20 km from Kribi on Lolodorf road, fr. 7 Aug. 1969, *Bos* 5154 (K, WAG); 20 km from Kribi, 3 km north of Lolodorf road, fr. 18 July 1969, *Bos* 5101 (K, WAG); **South West Region**, Ekumbe Mufako, fl., May 1994, *Ndam* 1198 (K, SCA, YA); Ekumbe Mufako, fl., May 1994, *Sonké* 1217 (K, SCA, YA); Ekumbe Mufako, fl., May 1994, *Thomas* 10121 (K, SCA, YA); Bipindi, fl., 1896, *Zenker* 1078 (K); Bipindi, fl. 1896, *Zenker* 1079 (K, P); Bipindi, fl. 1904, *Zenker* 2680 (BM, K).

## HABITAT

*Rinorea amietii* is known from littoral lowland evergreen rain forests. The altitude varies from 0 to 300 m.

### CONSERVATION STATUS

The littoral lowland evergreen rain forests of Cameroon are rich in *Rinorea* species. Many of the species are narrow endemics and so are vulnerable to extinction from forest clearance. Littoral forest habitat is highly threatened due to its ability to host industrial agriculture and because it is a target for logging activity. Many industrial farms are already located there, cultivating crops such as banana (*Musa spp*.), palm oil (*Elaeis guineensis*), rubber (*Hevea brasiliensis*). Due to its proximity to the coastal ports and the density of roads, littoral forest habitat is in demand for clearance for the expansion of these industries.

Currently eight locations are recorded for *Rinorea amietii* (see the 23 specimens studied), after extensive surveys in Cameroon (see under *Rinorea villiersii*). The extent of occurrence is estimated as 9,545 km^2^, and the area of occupation as 60 km^2^. Given the threats above, we assess *Rinorea amietii* as Vulnerable VU B1+2ab(iii)

### ETYMOLOGY

The specific epithet commemorates Professor Jean Amiet, lecturer at University of Yaoundé. He studied the relationship between butterflies and plants, especially that between *Cymothoe* (the admirals or gliders with about 75 species in Africa) and the genus *Rinorea* (Amiet 1997; Amiet 2000), and inspired the first author in his life’s taxonomic research on the genus *Rinorea* in Africa (Amiet & Achoundong 1996). He is commemorated by several other species such as *Afrothismia amietii* Cheek (Thismiaceae, Cheek 2003). Amiet, an entomologist, came to understand in the 1980s that the *Cymothoe* butterflies he studied were often monophagous, the larvae of some species eating only one species of *Rinorea* (see introduction). However, at that point, these different species of *Rinorea* were not recognised by taxonomists because of the superficial similarity of many other species. To address this difficulty he recruited the first author!

> **3. *Rinorea dewildei*** Achound., sp. nov

Type: Cameroon, Central Region, 35 km NW d’ Eséka, fl. 23 March 1964, *W*.*J*.*J*.*O. de Wilde* 2218 (holotype WAG; isotypes K001381925, P, YA),

*Rinorea dewildei* ined. Achoundong (1996: 544), Amiet & Achoundong (1996: 310); Onana (2011: 150).

*Shrub or treelet* up to 5 m high. Young twigs puberulent. Leaves papery, lamina elliptic, 7-18 × 2-7 cm, apex broadly acuminate, base cuneate to obtuse, margin serrate, of lateral nerves 6-8 pairs, 3-4 of which are more prominent than the others, Stipules triangular 6 × 2 mm, mucronate, abaxial midrib pronounced, pubescent, margin ciliolate. Petiole 5-15 mm long, pubescent. *Inflorescence* terminal, glabrous, corymbose, 3.5-4 cm. long; bracts triangular, up to 1 × 1.5 mm, ciliolate. *Flowers* yellow, 4 × 2 mm; sepals yellow, triangular, 2.5 × 1.2 mm, ciliolate. Petals yellow, ovate, 4 × 1 mm, ciliolate; 3-nerved. Androecium 2-3 mm long; staminal tube urceolate, 1.5 mm long; margin free, undulate, with a dorsal slit; staminal filaments inserted on the inner face of the staminal tube; filament of the dorsal stamen inserted near the distal edge of the staminal tube, on the dorsal slit; anthers 0.5 mm long; connective appendage red, ovate 0.5-1 mm long; thecal appendage bifid. Ovary ovoid, glabrous, 1 mm long; style 2 mm long, expanded at the summit. *Fruit* white, ovoid, tuberculate, 1.5 (−2) x 1 (−1.5) cm. *Seeds* tetrahedric, 0.5 cm x 0.5 cm. Fig 3.

**Fig. 3.**
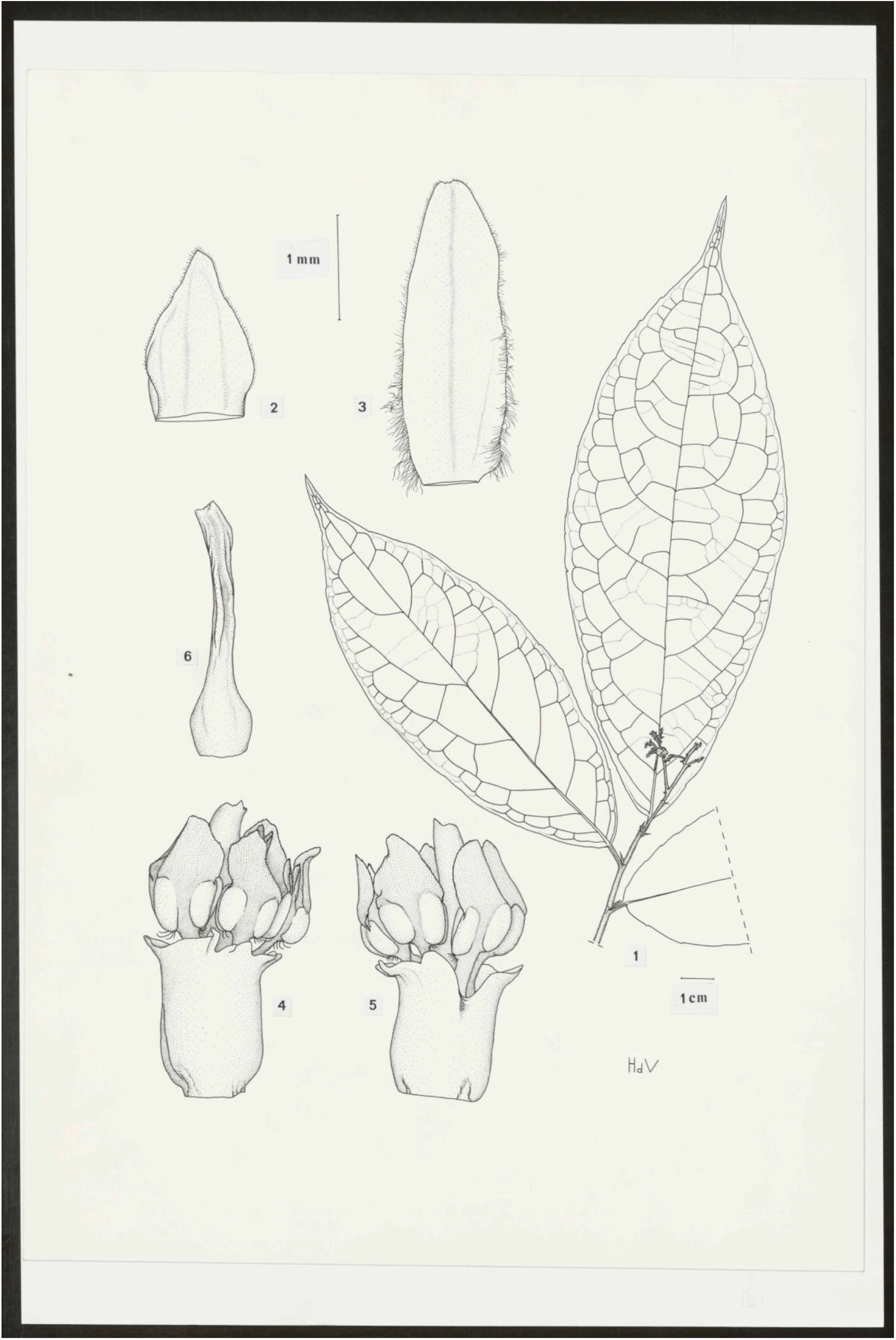
*Rinorea dewildei* **1** habit, flowering stem; **2** sepal; **3** petal; **4 & 5** staminal tube, side views; **6** pistil. All drawn from *W*.*J*.*J. O. de Wilde & B*.*E*.*E. de Wilde-Duyfjes* 2218 (WAG). by J.M. (HANS) DE VRIES

### RECOGNITION

In *Rinorea dewildei* the lamina is distinctive in that 3-4 of the lateral nerves are more prominent than the others. *Rinorea cerasifolia* Brandt shows the same character. The species are distinguished as followed:

> Petiole up to 5 cm long, tertiary venation of lamina sparse and inconspicuous; sepals ovate, as long as broad; all staminal filaments inserted on the inner face of the staminal tube ............................................................................... *Rinorea cerasifolia*
>
> Petiole 0.5-1.5 cm long; tertiary nervation of lamina dense and prominent; sepals triangular, twice as long as broad; one of the staminal filaments inserted on the slit of staminal tube margin, the others on the inner face .................... *Rinorea dewildei*

### DISTRIBUTION

Cameroon. *Rinorea dewildei* is found in the Central Region, Littoral Region and South Region (Map 1).

### SPECIMENS STUDIED. CAMEROON

**Littoral Region**, border of Nyong river at Dehane, fl., 1 March 1990, *Achoundong* 1623 (K, P, YA); ibid., *Achoundong* 2105 (YA); “Carriere Ducam Duclair”, 20 km north of Edea-Douala road, fl., 20 Feb. 2002, *Achoundong* 2197 (YA); 12 km Edea road, fl., 23 March 1994, *Achoundong* 2339 ED (P, YA); Edea area, fl., May 1994 *Achoundong* 17 ED (YA); **Central Region**, Makak, littoral forest, 100 km west of Yaoundé, young fr. Oct. 1938, *Jacques-Felix* 2288, (P); 35 km NW of Eséka, 23 Mar. 1964, *W*.*J*.*J*.*O. de Wilde* 2218 (holotype WAG; isotypes K, P, YA). 6 km NE of Edea, forest at Eding Island, 3°48.97’N, 10° 08.17’E, imm.fr., 7 May 2006, *van Velzen* 43 (WAG0416447). **South Region**. Bibabimvoto, Dipikar Island, in the Campo area along transect T2; fr., 19 July 2000, *Tchouto* T2X195 (WAG); Mabiogo, Dipikar island, in the Campo area along transect T1, fr., 20 Sep. 2000 *Tchouto* T1X144 (WAG); Bimbabivoto, Forest along transect T2 in Campo Maan area, fr., 18 July 2000 *Tchouto* 2951 (WAG).

### HABITAT

*Rinorea dewildei* occurs in littoral lowland evergreen rain forest. The altitude varies from 0 to 700 m. This species appears common in many areas of Cameroonian littoral forest but despite this has surprisingly few collections.

### CONSERVATION STATUS

According to Tchouto, *Rinorea dewildei* is common in the Campo-Ma’an National Park. Consequently, the species is well protected within the park. However, at locations outside the park, such as at Makak, Edea and Eseka, the habitat of this species is being cleared for agriculture and urbanization.

Currently, after extensive surveys in Cameroon (see under *Rinorea villiersii*), six locations are recorded for *Rinorea dewildei* (see specimens above), the extent of occurrence is estimated as 12,430 km^2^, and the area of occupation as 36 km^2^. Given the threats above, we assess *Rinorea dewildei* as VU B1+2ab(iii).

### ETYMOLOGY

The specific epithet commemorates Dr W.J.J.E. de Wilde. He and his wife, Brigitte de Wilde-Duyfjes collected the type specimen. Together, from their base at N’Kolbisson in Yaoundé in the 1960s, they collected numerous excellent botanical herbarium specimens in large sets through many parts of Cameroon including the littoral areas where *Rinorea dewildei* still grows. At this time their institute, Herbarium Vadense (WAG), then at Wageningen, Netherlands, was highly active in conducting botanical surveys in poorly studied but species-diverse areas of tropical Africa such as Cameroon and Gabon. Sixty years later, we are still naming new species to science resulting from their efforts, such as this.

### NOTES

One anomalous specimen from Gabon (Moyen-Ogooue, Camp Mboumi base, river bank, fr, fl. 20 Aug. 1999, *Issembe* 191 (K, LBV, WAG)), shares many similarities with *Rinorea dewildei*, and was initially identified as this species, but is here excluded. It may represent a further undescribed new species to science.

> ***4. Rinorea faurei*** *Achound*. sp. nov. Type: Cameroon, West Region, Santchou Forest Reserve, fl. *Achoundong* 1936 (holotype YA, isotypes P, WAG).

*Rinorea faurei* ined. Achoundong (1996: 544), Achoundong (1997: 255); Onana (2011: 150)

*Shrub or treelet* 1.5 – 2.5 (– 5) m high, stems and leaves glabrous. *Leaves* subcoriaceous, elliptic to slightly ovate, 12 – 21 × 4.5 – 6 (–9.5) cm, acuminate, base cuneate to obtuse, margin sightly serrate to subentire; lateral nerves 8 – 12 on each side of the midrib. Petiole 1 – 2.5 (– 5) cm long. Stipules triangular, 6 × 2 mm, mucronate, midrib abaxially pubescent, margin ciliolate. *Inflorescences* terminal or subterminal, thyrsiform, glabrous c.4 × 1.5 cm; bracts triangular, 1 × 1.5 mm, apiculate, abaxial midrib and margin ciliolate. *Flowers* white, zygomorphic, 4.5--6 mm long. Sepals 5, green, triangular to elliptic 1 – 1.3 × 3 × 1 – 1.5 mm. Petals white to yellow, elliptic, flat, 1.5 – 3 × 1 – 1.5 mm, apex rounded, margin ciliate. Lower petal broader than the upper and lateral petals, slightly constricted towards the apex, inner surface hairy near the apex. Androecium 3 mm long, staminal tube 0.3 – 1 mm long, margin free, with a V-shaped dorsal incision; filaments inserted on the inner face of the tube; anthers of the ventral side (face), fully exserted from the tube with free space between the tube summit and the lower part of the anthers; anthers 5, each 1 mm long, connective appendage red, ovate, 1 – 1.3 mm long, apex rounded, base decurrent on the upper part of the anthers. Ovary 3-locular, yellow, pyriform, 7 × 1 mm, style straight, cylindrical, white, 2 mm long, slightly constricted at the apex, stigma capitate. *Fruit* white, broader than long, 2 × 3 cm, strongly 3-lobed. *Seeds* six, white, acutely angled, dorsal face convex, ventral face flat c. 5 × 5 mm. Fig. 4

**Fig. 4.**
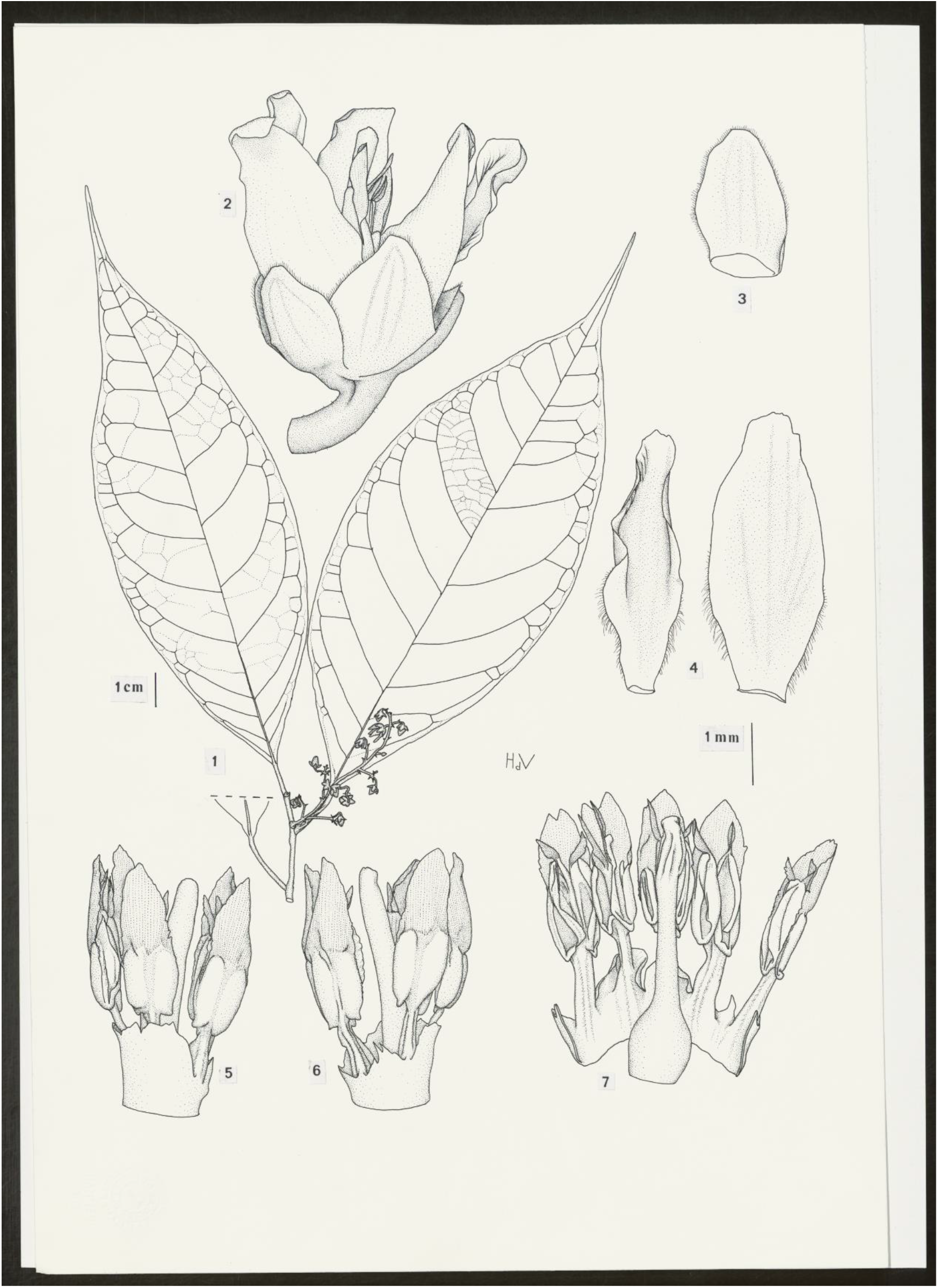
*Rinorea faurei* **1** habit, flowering stem; **2** flower, side view; **3** sepal; **4** petals, large (left) and small (right); **5 & 6** staminal tube and pistil, side views; **7** staminal tube and pistil, opened to show inner surface. All drawn from *Achoundong* 1936 (WAG) by J.M. (HANS) DE VRIES

## RECOGNITION

In its habit and its thyrsoid inflorescence *Rinorea faurei* is close to two other *Rinorea species* of the forest understory: *Rinorea sinuata* and *Rinorea thomasii*. The three species differ as follows:

1. Lamina oblong, shortly acuminate ........................................... *Rinorea thomasii*
1. Lamina elliptic or ovate to obovate, long-acuminate ........................................... 2
2. Inflorescence a dome-shaped, compact cyme ............................ *Rinorea sinuata*
2. Inflorescence an elongated, lax cyme, (flowers dispersed) ........... *Rinorea faurei*

### DISTRIBUTION

Cameroon. *Rinorea faurei* is endemic to the Santchou area in West Region (Map 1).

### SPECIMENS STUDIED. CAMEROON

**West Region**. Santchou Forest Reserve, fl., 17 March 1991, *Achoundong* 1786 (P, WAG, YA); Ibid., fl., fr. 19 May 1992 *Achoundong* 1925 (YA); ibid. *Achoundong* 1936 (holotype YA; isotypes P, WAG).

### HABITAT

*Rinorea faurei* occurs in understory of evergreen rain forest in the Santchou Plain at the foot of the Dschang Plateau. The estimated elevation above sea level is 770 m. The area of forest is more than 7000 ha. Santchou Forest Reserve is also the home of *Rinorea dentata, Rinorea oblongifolia, Rinorea yaundensis* and *Rinorea batesii* (Achoundong *pers. obs*. 1991-1992).

### CONSERVATION STATUS

Santchou Forest Reserve has official protection statutus. However, lack of concrete action on the ground means a strong risk of the reserve being invaded by farmers since the Santchou Plain is intensively cultivated. Currently, after extensive surveys in Cameroon (see under *Rinorea villiersii* above), only a single location in the sense of IUCN (2012) is recorded for *Rinorea faurei* (see specimens above), the extent of occurrence and the area of occupation is estimated as 70 km^2^. Given the threats above, we assess *Rinorea faurei* as Critically Endangered, CR B1+2ab(iii).

### ETYMOLOGY

The specific epithet honours Jean-Jacques Faure, former Head of the Forestry High School in Cameroon. His contribution to the protection of the environment of Central Africa has been important.

## Discussion

More collection is needed of *Rinorea* in Cameroon. The more widespread species such as *Rinorea dewildei* (see note there) have surprisingly few collections, even though the first author has noted they can be quite common within their range. It appears that many collectors, avoid targeting the genus in their botanical inventory work perhaps doubting that one species differs from another and seeking to avoid duplicating collections of the same taxon. The first author, in contrast, targeted the genus in his field surveys, accounting for his collections being the majority of those cited in this paper. This demonstrates how under-collected these Cameroon littoral forests are generally and confirms that more herbarium specimens should be collected to determine which species are present in these forests before they are lost.

**Map 1.**
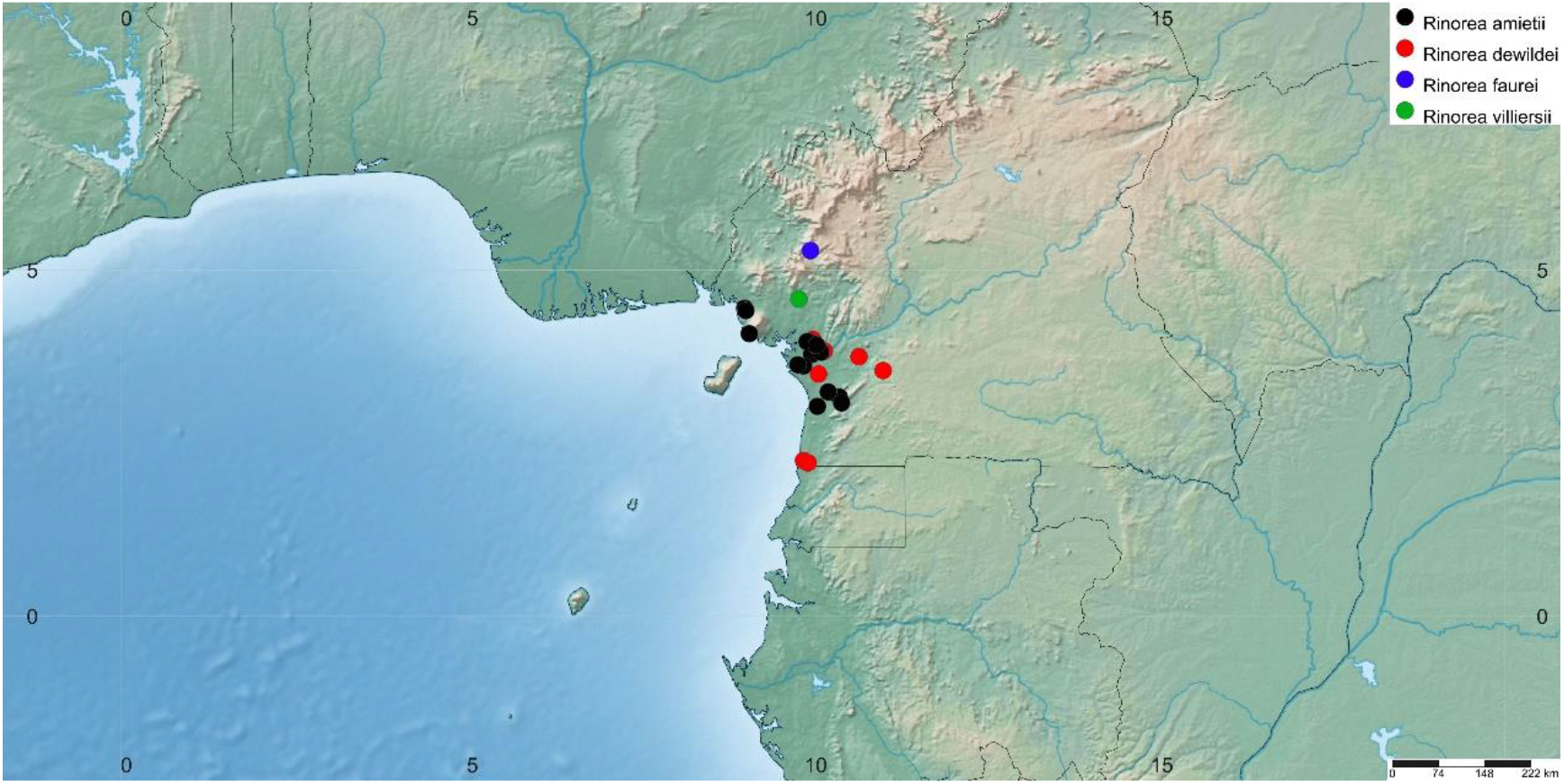
Global distributions of *Rinorea villiersii, R. amietii, R. dewildei* & *R. faurei*. Prepared with www.simplemappr.net

About 2000 species of vascular plant have been described as new to science each year for the last decade or more. Until species are known to science, they cannot be assessed for their conservation status and the possibility of protecting them is reduced (Cheek *et al*. 2020). To maximise the survival prospects of range-restricted species there is an urgent need not only to document them formally in compliance with the requirements of the relevant nomenclatural code (Turland *et al*. 2018), but also to formally assess the species for their extinction risk, applying the criteria of a recognised system, of which the IUCN Red List of Threatened Species is the most widely accepted (Bachman *et al*. 2019). Despite rapid increases over recent years in numbers of plant species represented by assessments on the Red List, the vast majority of plant species still lack such assessments (Nic Lughadha *et al*. 2020).

Increasing representation of plants on the Red List is imperative but documenting threatened species in the level of detail required for inclusion on the Red List is resource intensive (Juffe-Bignoli *et al*. 2016) and securing the required peer-review for completed assessments to be accepted for the Red List can be slow and problematic (Bachman *et al*. 2019). Alternative approaches to evaluating extinction risk, including automated or semi-automated methods involving digitally available information and artificial intelligence, have been applied to plants in recent years (e.g., Darrah *et al*. 2017, Nic Lughadha *et al*. 2019, Zizka *et al*. 2020), often with demonstrable success in correctly predicting the threat status of species previously assessed for the Red List. However, caution is needed in considering such automated assessment approaches for adoption as conservation tools, not only for a variety of technical reasons (Walker *et al*. 2020) but also because they lack the power that an IUCN Red List assessment has to help change the fate of a species.

As a global standard, the IUCN Red List not only provides stable and evidence-based methods for biodiversity scientists to assess and document the extinction risk of individual species, thus facilitating comparison of the extinction risk of taxonomic groups and areas worldwide (e.g., Nic Lughadha *et al*. 2020; Mair *et al*. 2021), it also supports the safeguarding and sustainability frameworks used by businesses and their major lenders (Bennun *et al*. 2018; Juffe-Bignoli *et al*. 2016). For example, clients of the International Finance Corporation (World Bank Group) are required to use the Red List to inform project risks and to refrain from activities leading to a net reduction in populations of species assessed on the Red List as Endangered (EN) or Critically Endangered (CR), over a reasonable timescale. Species whose likely extinction has been avoided as a result of these rules include *Stylochaeton pilosum* Bogner, *Marsdenia exellii* C.Norman, *Raphionacme caerulea* E.A.Bruce (all EN), and *Tarenna hutchinsonii* Bremek. (CR) (Couch *et al*. 2014; Couch *et al*. 2019).

National governments and leaders also recognise the importance of species assessed as threatened by on the Red List, as demonstrated recently in Cameroon when in part due to the high number of plant species on the Red List (Lovell 2020), a logging concession was revoked for the Ebo forest (Kew Science News 2020).

Thus, while the Conservation Status statements included in the species treatments in the present paper suffice to highlight the threatened status of these species to the taxonomic botany community, completion and review of full assessments must follow so that they can be published on the IUCN Red List. Their assessments on the Red List will facilitate their inclusion in larger scale studies, including future global extinction risk estimates and conservation prioritisation exercises, but, most importantly, it will enable not scientists, local communities, NGOs and national authorities to promote and take action to safeguard them.

Documented extinctions of plant species are increasing (Humphreys *et al*. 2019) and recent estimates suggest that as many as two fifths of the world’s plant species are now threatened with extinction (Nic Lughadha *et al*. 2020). In Cameroon, *Oxygyne triandra* Schltr. and *Afrothismia pachyantha* Schltr. of South West Region, Cameroon are now known to be globally extinct (Cheek & Williams 1999, Cheek *et al*. 2018a, Cheek *et al*., 2019). In some cases, Cameroon species appear to have become extinct even before they are known to science, such as *Vepris bali* Cheek (Cheek *et al*. 2018b), Most of the 815 Cameroonian species in the Red Data Book for the plants of Cameroon are threatened with extinction due to habitat clearance or degradation, especially of forest for small-holder and plantation agriculture following logging (Onana & Cheek, 2011). Efforts are now being made to delimit the highest priority areas in Cameroon for plant conservation as Tropical Important Plant Areas (TIPAs) using the revised IPA criteria set out in Darbyshire *et al*. (2017). This is intended to help avoid the global extinction of additional endemic species such as the *Rinorea* species published in this paper which it is intended will be included in the future proposed TIPAs.

## Acknowledgements

The first author thanks IRAD-National Herbarium of Cameroon (YA) for support in his retirement that has enabled him to continue and finalise his taxonomic research on Violaceae of Cameroon for the Flore du Cameroun. He also thanks colleagues at Wageningen for retrieving and transmitting the excellent figures that illustrate this paper, mainly drawn by J.M. (Hans) de Vries, with valuable contributions by Jean Michel Onana. This paper was completed as part of the Cameroon TIPAs (Tropical Important Plant Areas) project at RBG, Kew, which is supported by Players of Peoples Postcode Lottery. Formal red listing of these four species, once they are formally published, will be supported by the John S. Cohen Foundation.

## References

Achoundong, G. (1996). Les Rinorea comme indicateurs des grands types forestiers du Cameroun. In: van der Maesen, L.J.G., van der Burgt, X.M. & Van Medenbach-de Rooy, J.M. (eds) The Biodiversity of African Plants. Kluwer Academic Publishers, Dordrecht, pp. 536–544. http://dx.doi.org/10.1007/978-94-009-0285-5_69

Achoundong, G. (1997). Rinorea du Cameroun, Systématique, Biologie, Ecologie, Phytogéographie. Thèse Université de Yaoundé I, Yaoundé.

Achoundong, G. (2000). Les Rinorea et l’étude des refuges forestiers en Afrique. In: Servant, M. & Servant-Vildary, S. (eds) Dynamique à long terme des écosystèmes forestiers intertropicaux. Mémoire UNESCO, Paris. pp 19–29

Achoundong, G. (2003) Novitates Gabonenses 45. Une nouvelle espèce de Rinorea (Violaceae) du Gabon. Adansonia n.s. 25: 211–215.

Achoundong, G. & Bakker, F. T. (2006). Deux nouvelles espèces de Rinorea, série Ilicifoliae (Violaceae) du Cameroun. Adansonia, sér. 3, 28 (1): 129–136.

Achoundong, G. & Bos, J.J. (1999). Novitates Gabonenses: 37. Espèces nouvelles de Rinorea (Violaceae) du Gabon. Adansonia 21 (1): 125–131

Achoundong, G. & Bos, J.J. (2001). Deux espèces nouvelles de Rinorea (Violaceae) du Congo et du Gabon. Adansonia 23 (1): 155–159.

Achoundong, G. & Cheek, M. (2003). Two new species of Rinorea (Violaceae) from western Cameroon. Kew Bulletin 58: 957–964. https://doi.org/10.2307/4111209

>Achoundong G. & Cheek, M. (2005). Two further new species of Rinorea (Violaceae) from Cameroon. Kew Bulletin 60 (4): 581–586

Achoundong, G. & Onana, J. (1998). Allexis zygomorpha (Violaceae): a new species from the Littoral Forest of Cameroon. Kew Bulletin 53 (4): 1009–1010. https://doi.org/10.2307/4118897

Amiet, J.-L. (1997) Spécialisation trophique et premières états chez les Cymothoe: implications taxonomiques (Lepidoptera, Nymphalidae). Bulletin de la Société entomologique de France 102: 15–29.

Amiet, J.-L. (2000) Les premiers états des Cymothoe: Morphologie et intérêt phylogénique (Lepidoptera, Nymphalidae). Bulletin de la Société entomologique de France 106: 349–390

Amiet, J. L. & Achoundong, G. (1996). Un exemple de speciation trophique chez les Lepidopteres: les Cymotoe camerounaises infeodees au Rinorea (Violacies) (Lepidoptera, Nymphalidae). Bull. Soc. Entom. Fr. 105(5): 449–466.

Bachman, S.P., Field, R., Reader, T., Raimondo, D., Donaldson, J., Schatz, G.E. and Lughadha, E.N. (2019). Progress, challenges and opportunities for Red Listing. Biological Conservation, 234: 45–55. https://doi.org/10.1016/j.biocon.2019.03.002

Bachman, S., Moat, J., Hill, A., de la Torre, J., Scott, B. (2011). Supporting Red List threat assessments with GeoCAT: geospatial conservation assessment tool. ZooKeys. 150: 117–126. https://doi.org/10.3897/zookeys.150.2109

Bakker, F. T., van Gemerden, B. S. & Achoundong, G. (2006). Molecular systematics of African Rinorea Aublet. (Violaceae). Pp. 33–44 in S. A. Ghazanfar and H. J. Beentje (eds.) Taxonomy and ecology of African plants, their conservation and sustainable use. Royal Botanic Gardens, Kew.

Bennun, L., Regan, E.C., Bird, J., van Bochove, J.W., Katariya, V., Livingstone, S., Mitchell, R., Savy, C., Starkey, M., Temple, H. and Pilgrim, J.D., 2018. The value of the IUCN Red List for business decision-making. Conservation Letters, 11(1), p.e12353.https://doi.org/10.1111/conl.12353

Chapman, J. & Chapman, H. (2001). The Forests of Taraba and Adamawa States, Nigeria an Ecological Account and Plant Species Checklist. University of Canterbury: Christchurch, New Zealand. pp. 221.

Cheek, M. (2017). Rinorea simoneae. The IUCN Red List of Threatened Species 2017: e.T110087540A110087542. https://dx.doi.org/10.2305/IUCN.UK.2017-3.RLTS.T110087540A110087542.en.

Cheek, M., Gosline, G., Onana, J-M. (2018b). Vepris bali (Rutaceae), a new critically endangered (possibly extinct) cloud forest tree species from Bali Ngemba, Cameroon. Willdenowia 48: 285–292. doi:https://doi.org/10.3372/wi.48.48207

Cheek, M., Harvey Y., Onana J-M. (2010). The Plants of Dom, Bamenda Highlands, Cameroon, A Conservation Checklist. Kew, Royal Botanic Gardens.

Cheek, M., Harvey, Y., Onana, J.-M. (2011). The Plants of Mefou Proposed National Park, Yaoundé, Cameroon, A Conservation Checklist. Kew, Royal Botanic Gardens.

Cheek, M., Mackinder, B. Gosline, G., Onana J.-M., Achoundong, G. (2001). The phytogeography and flora of western Cameroon and the Cross River-Sanaga River interval. Systematics and Geography of Plants 71: 1097–1100. https://doi.org/10.2307/3668742

Cheek, M., Nic Lughadha, E., Kirk, P., Lindon, H., Carretero, J., Looney, B.,Douglas, B., Haelewaters, D., Gaya, E., Llewellyn, T., Ainsworth, M.,Gafforov, Y., Hyde, K., Crous, P., Hughes, M., Walker, B.E., Forzza, R.C., Wong, K.M., Niskanen, T. (2020). New scientific discoveries: plants and fungi. Plants, People Planet 2:371–388. https://doi.org/10.1002/ppp3.10148

Cheek, M., Onana, J-M., Pollard, B.J. (2000). The Plants of Mount Oku and the Ijim Ridge, Cameroon, a Conservation Checklist. Kew, Royal Botanic Gardens.

Cheek, M., Pollard, B.J., Darbyshire, I., Onana, J-M., Wild, C. (2004). The Plants of Kupe, Mwanenguba and the Bakossi Mountains, Cameroon: A Conservation Checklist. Kew, Royal Botanic Gardens.

Cheek, M., Tsukaya, H., Rudall, P.J., Suetsugu, K. (2018a). Taxonomic monograph of Oxygyne (Thismiaceae), rare achlorophyllous mycoheterotrophs with strongly disjunct distribution. PeerJ, 6, e4828. https://doi.org/10.7717/peerj.4828

Cheek, M. & Williams, S. (1999). A Review of African Saprophytic Flowering Plants. In: Timberlake, Kativu eds. African Plants. Biodiversity, Taxonomy & Uses. Proceedings of the 15th AETFAT Congress at Harare. Zimbabwe, 39–49.

Couch, C., Molmou, D., Camara, B., Cheek, M., Merklinger, F., Davies, L., Harvey, Y., Lopez Poveda, L. and Redstone, S., 2014. Conservation of threatened Guinean inselberg species. In XXth AETFAT Congress, South Africa, 2014 [Abstract 96]. Scripta Botanica Belgica (52 : 96).

Couch, C., Cheek, M., Haba, P., Molmou, D., Williams, J., Magassouba, S., Doumbouya, S. and Diallo, M.Y., 2019. Threatened Habitats and Tropical Important Plant Areas (TIPAs) of Guinea, West Africa. Kew: Royal Botanic Gardens, Kew.

Darbyshire, I. & Cheek, M. (2004a). Rinorea fausteana. The IUCN Red List of Threatened Species 2004: e.T46185A11034817. https://dx.doi.org/10.2305/IUCN.UK.2004.RLTS.T46185A110 34817.en.

Darbyshire, I. & Cheek, M. 2004b. Rinorea thomasii. The IUCN Red List of Threatened Species 2004: e.T46186A11034916. https://dx.doi.org/10.2305/IUCN.UK.2004.RLTS.T46186A110 34916.en.

Darbyshire, I., Anderson, S., Asatryan, A., Byfield, A., Cheek, M., Clubbe, C., Ghrabi, Z., Harris, T., Heatubun, C. D., Kalema, J., Magassouba, S., McCarthy, B., Milliken, W., Montmollin, B. de, Nic Lughadha, E., Onana, J. M., Saidou, D., Sarbu, A., Shrestha, K. & Radford, E. A. (2017). Important Plant Areas: revised selection criteria for a global approach to plant conservation. Biodiversity Conservation 26: 1767–1800. https://doi.org/10.1007/s10531-017-1336-6

Darrah, S.E., Bland, L.M., Bachman, S.P., Clubbe, C.P. &Trias-Blasi, A. (2017). Using coarse-scale species distribution data to predict extinction risk in plants. Diversity and Distributions,23:435–447. https://doi.org/10.1111/ddi.12532

Harvey, Y., Pollard, B.J., Darbyshire, I., Onana, J-M., Cheek, M. (2004). The Plants of Bali Ngemba Forest Reserve, Cameroon. A Conservation Checklist. Kew, Royal Botanic Gardens.

Harvey, Y.H., Tchiengue, B., Cheek, M. (2010). The plants of the Lebialem Highlands, a conservation checklist. Kew, Royal Botanic Gardens.

Humphreys, A.M., Govaerts, R., Ficinski, S.Z., Lughadha, E.N. and Vorontsova, M.S. (2019) Global dataset shows geography and life form predict modern plant extinction and rediscovery. Nature Ecology & Evolution 3.7: 1043–1047. https://doi.org/10.1038/s41559-019-0906-2

IPNI (continuously updated). The International Plant Names Index. Available from: http://ipni.org/ [accessed Mar. 2018].

IUCN (2012). IUCN red list categories: Version 3.1. Gland, Switzerland and Cambridge, U.K., IUCN Species Survival Commission.

Juffe-Bignoli, D., Brooks, T.M., Butchart, S.H., Jenkins, R.B., Boe, K., Hoffmann, M., Angulo, A., Bachman, S., Böhm, M., Brummitt, N. & Carpenter, K.E. (2016). Assessing the cost of global biodiversity and conservation knowledge. PLoS One 11(8) https://doi.org/10.1371/journal.pone.0160640

Kew Science News (2020). Ebo Forest Logging Plans Suspended. https://www.kew.org/read-and-watch/ebo-forest-logging-suspended (accessed 5 May 2021).

Lovell, R. (2020). Timber ! The threat to Cameroon’s Ebo Forest. https://www.kew.org/read-and-watch/ebo-forest-cameroon (accessed 5 May 2021).

Mair, L., Bennun, L.A., Brooks, T.M., Butchart, S.H., Bolam, F.C., Burgess, N.D., Ekstrom, J.M., Milner-Gulland, E.J., Hoffmann, M., Ma, K. and Macfarlane, N.B., (2021). A metric for spatially explicit contributions to science-based species argets. Nature Ecology & Evolution, pp.1–8. https://doi.org/10.1038/s41559-021-01432-0

Nic Lughadha, E., Walker, B.E., Canteiro, C., Chadburn, H., Davis, A.P., Hargreaves, S., Lucas, E.J., Schuiteman, A., Williams, E., Bachman, S.P. and Baines, D.(2019). The use and misuse of herbarium specimens in evaluating plant extinction risks. Philosophical transactions of the Royal Society B, 374(1763), p.20170402. https://doi.org/10.1098/rstb.2017.0402

Nic Lughadha, E., Bachman, S.P., Leão, T.C., Forest, F., Halley, J.M., Moat, J., Acedo, C., Bacon, K.L., Brewer, R.F., Gâteblé, G. and Gonçalves, S.C. (2020). Extinction risk and threats to plants and fungi. Plants, People, Planet, 2(5), pp.389-408. https://doi.org/10.1098/rstb.2017.0402

Onana, J.-M. (2011). The vascular plants of Cameroon, a taxonomic checklist with IUCN Assessments. Kew, Royal Botanic Gardens.

Onana, J.-M., Cheek, M. (2011). Red data book of the flowering plants of Cameroon, IUCN global assessments. Kew, Royal Botanic Gardens.

Sosef, M.S.M., Wieringa, J.J., Jongkind, C.C.H., Achoundong, G., Azizet Issembé, Y., Bedigian, D., Van Den Berg, R.G., Breteler, F.J., Cheek, M., Degreef, J. (2006). Checklist of Gabonese Vascular Plants. Scripta Botanica Belgica 35. National Botanic Garden of Belgium.

Turland, N. J., Wiersema, J. H., Barrie, F. R., Greuter, W., Hawksworth, D. L., Herendeen, P. S., Knapp, S., Kusber, W.-H., Li, D.-Z., Marhold, K., May, T. W., McNeill, J., Monro, A. M., Prado, J., Price, M. J. & Smith, G. F. (eds.) (2018). International Code of Nomenclature for algae, fungi, and plants (Shenzhen Code) adopted by the Nineteenth International Botanical Congress Shenzhen, China, July 2017. Regnum Vegetabile 159. Glashütten: Koeltz Botanical Books. DOI https://doi.org/10.12705/Code.2018

van Velzen, R., & Wieringa, J. J. (2014). Rinorea calcicola (Violaceae), an endangered new species from south-eastern Gabon. Phytotaxa, 167 (3): 267–275. https://doi.org/10.11646/phytotaxa.167.3.5

van Velzen, R., Wahlert, G.A., Sosef, M.S., Onstein, R.E., Bakker, F.T. (2015). Phylogenetics of African Rinorea (Violaceae): Elucidating Infrageneric Relationships using Plastid and Nuclear DNA Sequences, Systematic Botany 40 (1). 174–184. https://doi.org/10.1600/036364415X686486

Wahlert, G. A. (2010). Phylogeny, biogeography, and a taxonomic revision of Rinorea (Violaceae) from Madagascar and the Comoro Islands. Ph.D. Thesis. Athens: Ohio University.

Wahlert, G. A. & Ballard, H.E. (2012). A phylogeny of Rinorea (Violaceae) inferred from plastid DNA sequences with an emphasis on the African and Malagasy species. Systematic Botany 37: 964–973. https://doi.org/10.3389/fpls.2020.00520

Walker, B.E., Leão, T.C., Bachman, S.P., Bolam, F.C. and Nic Lughadha, E. (2020). Caution needed when predicting species threat status for conservation prioritization on a global scale. Frontiers in plant science 11:520. https://doi.org/10.3389/fpls.2020.00520

Zizka, A., Silvestro, D., Vitt, P. and Knight, T.M. (2020). Automated conservation assessment of the orchid family with deep learning. Conservation Biology. https://doi.org/10.1111/cobi.13616

Thiers, B. (continuously updated). Index Herbariorum: A global directory of public herbaria and associated staff. New York Botanical Garden’s Virtual Herbarium. [continuously updated]. Available from: http://sweetgum.nybg.org/ih/ [accessed: March 2021].

